# PickPocket: Pocket binding prediction for specific ligand families using neural networks

**DOI:** 10.1101/2020.04.15.042655

**Authors:** Benjamin Viart, Claudio Lorenzi, María Moriel-Carretero, Sofia Kossida

## Abstract

Most of the protein biological functions occur through contacts with other proteins or ligands. The residues that constitute the contact surface of a ligand-binding pocket are usually located far away within its sequence. Therefore, the identification of such motifs is more challenging than the linear protein domains. To discover new binding sites, we developed a tool called PickPocket that focuses on a small set of user-defined ligands and uses neural networks to train a ligand-binding prediction model. We tested PickPocket on fatty acid-like ligands due to their structural similarities and their under-representation in the ligand-pocket binding literature.

Our results show that for fatty acid-like molecules, pocket descriptors and secondary structures are enough to obtain predictions with accuracy >90% using a dataset of 1740 manually curated ligand-binding pockets. The trained model could also successfully predict the ligand-binding pockets using unseen structural data of two recently reported fatty acid-binding proteins. We think that the PickPocket tool can help to discover new protein functions by investigating the binding sites of specific ligand families. The source code and all datasets contained in this work are freely available at https://github.com/benjaminviart/PickPocket.

**Author Summary:** Most of the protein biological functions are defined by its interactions with other proteins or ligands. The cavity of the protein structure that receives a ligand, also called a pocket, is made of residues that are usually located far away within its sequence. Therefore understanding the omplementarity of pocket and ligand is a real challenge. To discover new binding sites, we developed a tool called PickPocket that focuses on a small set of user-defined ligands to train a prediction model. Our results show that for fatty acid-like molecules, pocket descriptors (such as volume, shape, hydrophobicity…) and secondary structures are enough to obtain predictions with accuracy >90% using a dataset of 1740 manually curated ligand-binding pockets. The trained model could also successfully predict the ligand-binding pockets using unseen structural data of two recently reported fatty acid-binding proteins. We think that the PickPocket tool can help to discover new protein functions by investigating the binding sites of specific ligand families.

## Introduction

One of the main tasks of bioinformatics is to associate biological roles to proteins using the always increasing biological data (1,2). To predict the function of a protein based on its sequence, computational methods look for sequence patterns in biological databases of known and already annotated proteins. Homology search (3,4), motif search (5) and functional domain search (6,7) are the most common methods, among the many available tools. Other strategies exploit different data types, such as gene expression (8) or even a combinatorial approach (9,10).

Most of the protein biological functions occur through interactions with other proteins or ligands (11). The residues making the protein contact surface and cavity shape are often located far away within the protein sequence. Therefore, the identification of such motifs is more difficult. Fortunately, the quantity of protein structures and models available in the Protein Data Bank (PDB) archive (12) has increased rapidly (13), providing abundant data for structural analyses. The study of ligands and cavities or pockets to which they bind is of particular interest, especially for drug discovery.

Different algorithms exist to compute the structure of ligand-binding pockets and to predict binding sites using geometric criteria, such as SURFNET (14), APROPOS (15), Q-SiteFinder (16), LIGSITEcsc (17), ConCavity (17,18), fpocket (19),DEPTH (20) or PDBinder (21) among others. Artificial intelligence also has been useful in this field with the development of algorithms for convolutional neural networks, such as FRSite (22), DeepSite (23) and DeepDrug3D (24) or of tools based on the random forest algorithm, such as P2Rank (25).

With the progressive increase in structure availability, the need to store and compare ligand-binding pocket data has led to the creation of dedicated databases, such as the Computed Atlas of Surface Topography of proteins (CASTp) (26,27) and the PoSSuM database (28). The collection of all pockets present in a single organism is called a pocketome and has its dedicated database: Pocketome (29). The extensive knowledge of all the pocket structures and their comparison are valuable for drug designers. For special needs, the tool PocketPipe can be used to analyze the pocketome of a single organism (30). Recently, comparative analyses of binding sites have gained momentum due to their capacity to reveal ligand-binding similarities among proteins, regardless of their evolution (31,32). As different proteins can evolve to bind to the same ligand type (33) the accurate classification of binding sites has become an important tool for designing drugs and predicting their possible side effects through unwanted binding (34,35).

One limitation of the available tools is that they are tailored for drug design or for the analysis of large pockets, thus excluding the possibility of executing other tasks, such as determining the specific ligand-pocket binding complementarity. Yet, to discover new binding sites, the reverse approach needs to be possible: to focus on a small set of specific ligands and to take into account their molecular and structural specificity.

In this work, we developed a tool called PickPocket to generate a dataset of pockets that interact with a specific user-defined ligand family. The workflow consists of different R (36) and python (36,37) programs organized using bash scripts. From the pocket dataset, we computed a descriptive matrix of all pockets that contains structural information on the cavity and the residue secondary structure. Then, we trained a neural network multilayer perceptron (38) to predict whether a cavity is a ‘true’ pocket (i.e. can interact with ligands of the family under study) or a ‘false’ pocket.

To test PickPocket, we decided to use fatty acid-like ligands due to their structural similarities and their under-representation in the ligand-pocket binding literature (39). The source code and all datasets contained in this work are freely available at https://github.com/benjaminviart/PickPocket.

## Results

The input data for the PickPocket tool consisted of 42 fatty acid-like ligands (Supplementary Information Table 1: Detail of ligands used as input for PickPocket), 301 structures containing one of the selected ligands and 242 structures containing no ligand. For each pocket, a descriptive matrix is computed that gathers 21 features including structural information of the cavity as well as secondary structure (Table 1). In order to ensure the quality of the training data, the pockets labels were manually checked using pymol (40). A few mis-annotations were detected and corrected. The final training matrix contained 339 ligand-binding (true) pockets and 1401 empty ones (false).

**Table 1:** Descriptors used to train the predictive model

PickPocket: Pocket binding prediction for specific ligand families using neural networks.
|  |  |
| --- | --- |
| Pocket score | Charge score |
| Druggability score | Local hydrophobic density score |
| Number of vertices | Number of apolar alphaspheres |
| Mean radius of alpha sphere | Proportion of apolar alpha sphere |
| Mean alpha sphere solvent accessibility | Solvent accessible surface |
| Mean B factor | Number of alpha helices |
| Hydrophobicity score | Number of coils |
| Polarity score | Number of strands |
| Volume score | Number of turns |
| Real volume | Number of bridges |
|  | Number of 310 helices |

**Table 2:** Prediction results. Prediction scores for the pockets of structure 6S2M and 6U1U. Score >0.5 are considered positives and colored green. Color corresponds to Figure 2. Complete prediction results can be found in supplementary information (SI Table 2: Full prediction results for 6S2M and 6U1U PDB structures).

PickPocket: Pocket binding prediction for specific ligand families using neural networks.
| Structure | Pocket color | Prediction Score |
| --- | --- | --- |
| 6S2M(A) | RED | 0.78 |
| 6S2M(A) | BLUE | 0.03 |
| 6S2M(A) | WHITE | 0.18 |
| 6S2M(A) | YELLOW | 0.2 |
| 6S2M(A) | PURPLE | 0.04 |
| 6U1U(B) | RED | 0.69 |
| 6U1U(B) | YELLOW | 0.0 |
| 6U1U(B) | PURPLE | 0.0 |
| 6U1U(B) | LIGHT GREEN | 0.19 |
| 6U1U(B) | GREEN | 0.0 |

We obtained the best results using a neural network multilayer perceptron classifier with an architecture of (15, 10, 5). To avoid overfitting, we trained the model using a 5-fold cross-validation. Furthermore, in order to reduce problems associated with unbalanced classes, we downsampled the largest groups according to the smallest one. The model displayed an Area Under the Curve (AUC) of 97.2% (ROC curve in Figure 1). The model accuracy was 93.4%.

**Figure 1:**
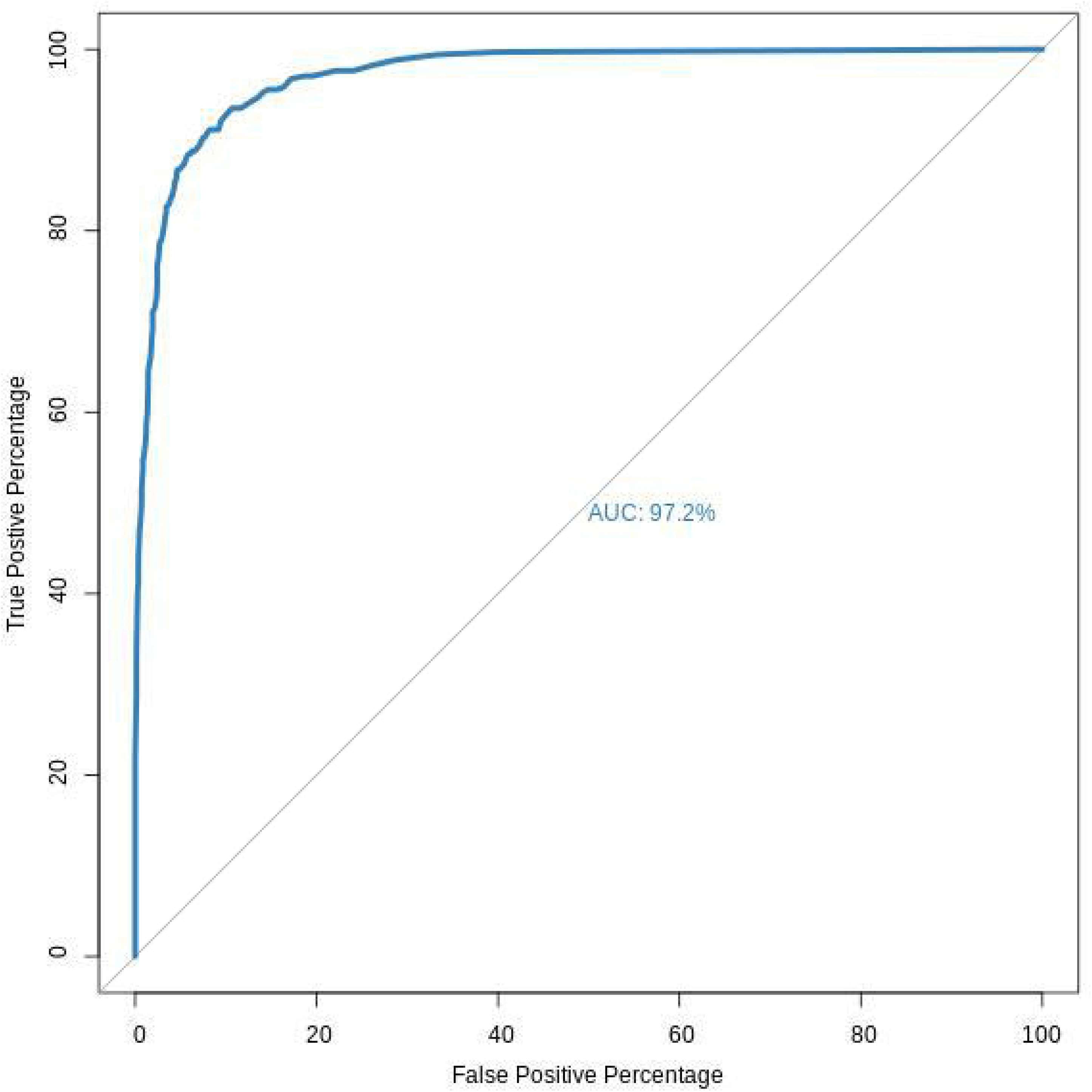
ROC curve for pocket prediction after 5-fold cross-validation. The Area Under the Curve (AUC) of the best model was 97.2%.

To demonstrate that PickPocket can predict ligand binding on unseen data, we selected the two most recent (at the time of writing) human protein structures containing fatty acid(s) from the PDB archive: perdeuterated human myelin protein P2 (PDBID=6S2M), which contains two possible fatty acids in the same pocket (vaccenic acid and palmitic acid), and human angiopoietin-like 4 (PDBID=6U1U), which contains palmitic acid. Pockets with a score ≥0.5 were considered positive and were colored in red, the others were in random colors. For both structures, PickPocket correctly identified the fatty acid binding cavity. Careful analysis of the 6S2M structure showed that the cavity fatty acid occupied two pockets (red and blue in Figure 2A). The red pocket, which is the deep part inside the protein and contained the carboxyl part, had a score of 0.78. The blue pocket, which is at the opening of the cavity and contained the fatty acid tail, had a score of 0.03. The fatty acid-binding cavities were large and, as illustrated in this case, fpocket tended to consider them as more than one pocket. As both pockets corresponded to the same cavity and the red pocket had a score well-above the threshold, we considered that PickPocket discovered the fatty acid-binding cavity of the structure. For 6U1U (Figure 2B), the red pocket, corresponding to the palmitic acid cavity, received a score of 0.69.

**Fig 2.**
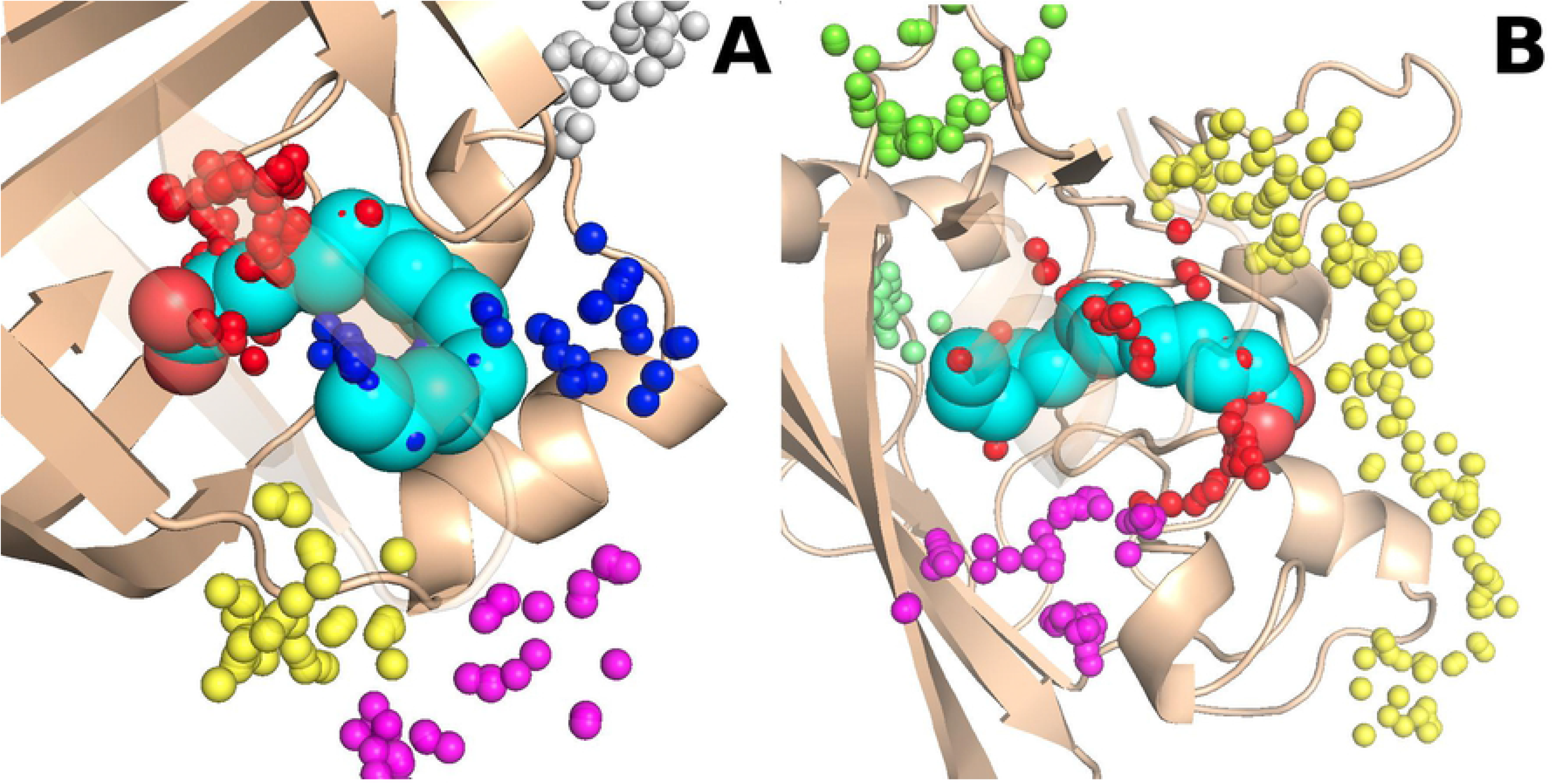
Prediction details for the recently published structures 6S2M. **(A) and 6U1U (B).** Each set of colored balls represents a pocket. Only the relevant parts of the structure are shown here. Some parts of the protein were set transparent to facilitate the visualization, and some negative pockets distant from the fatty acid were hidden to help visibility. The red pockets are correctly predicted, while the blue pocket, which corresponds to the palmitic acid tail, was incorrectly categorized as false.

## Discussion and perspectives

Thanks to the increased availability of structural data and the improvement of protein 3D modeling, new pocket-based methodologies can now be developed to help protein function discovery. Our results show that combining pocket descriptors and the residue secondary structure is sufficient to train a model and to predict the pockets for specific ligand families with high accuracy, including when using unseen data. We believe that careful analysis of protein ligand families and their corresponding binding pockets coupled with the high prediction capacity of neural networks is the way forward to close the gap between protein sequences and their functions. PickPocket aims to simplify the procedure for building a ligand family dataset and train a model to recognize the corresponding cavity. The ligand selection and the corresponding structure still need to be done manually, but these steps are made easier by the PDB ligand research tool. Other databases, such as PoSSuM (28), may be used to create the dataset.

PickPocket automatic labelling of ‘true’ and ‘false’ pockets is very fast, but still needs manual checking because sometimes it makes mistakes when pockets are very close to each other. The training data quality is extremely important for good predictions. Therefore, we strongly recommend users to manually check the generated dataset. Looking in detail at the results for the 6S2M structure, fpocket considered that it had two different pockets because the fatty acid binding cavity is very deep and narrow. This can be corrected by changing the alpha-sphere radius or the maximum distance between pockets. One of the challenges we faced was to tune the fpocket parameters in order to have a big enough pocket size without merging different cavities. PickPocket easy tuning of these parameters allows users to adapt the input to the ligand specificity.

In order to cluster ligand protein complexes, Deepdrug3D and FRsite use an atom-based voxelization. This step also allows generating a compatible input for convolutional neural networks. On the other hand, our methodology uses matrix properties that are faster to generate, but contain less information. We also chose to use the fpocket software, although DeepSite is more accurate against the sc-PDB database of binding sites (40). However, fpocket is fast, and pocket descriptors data is easily retrieved from output files.

PickPocket can help to discover new protein functions by investigating the binding sites of a specific ligand family. The results we obtained prove that for fatty acid-like molecules, pocket descriptors and secondary structure are enough to obtain predictions with >90% accuracy. Thanks to its high prediction accuracy, PickPocket can be used as a tool for *in silico* screens, and should boost novel research.

## Material and Methods

PickPocket methodology can be divided into five steps (Figure 3). First a selection of ligands and structure, second the combination of fpocket and Stride to generate the descriptive matrice of the pockets, third the selection of ‘true’ pocket (pocket that binds the selected ligands), fourth training the desired model and last computing the prediction on the full PDB pocket dataset.

**Figure 3:**
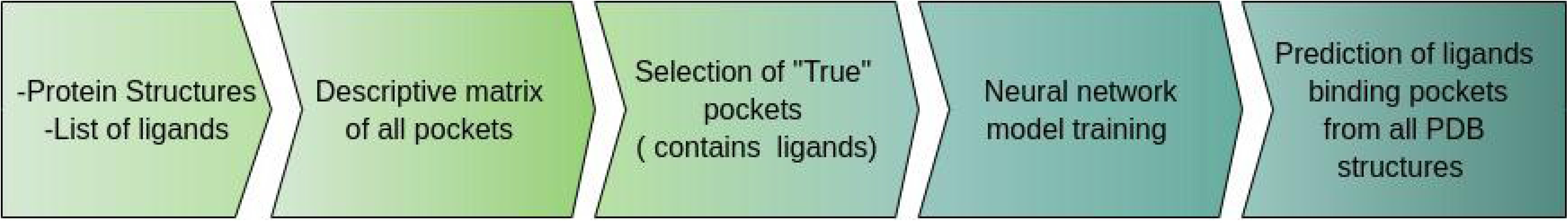
methodology workflow. Five steps of the PickPocket methodology for specific ligands binding prediction.

To select a set of fatty acid-like ligands present in the PDB we used the ligand search tool. We selected ligands having this SMILES ‘CCCCCCCCCCCCCCCCCCCCCCCCCCCCCCC(O)=O’ as superstructure and a molecular weight superior to 100.0 g/mol.

This resulted in a list of mostly fatty acids and other molecules composed of aliphatic chains including some with alcohols.From the structures containing our ligands we only selected the one with human proteins, resolved using X-ray technique and excluding DNA and RNA molecules. A representative subset at 70% sequence identity was then used as the input. A set of random structures, using the same criteria (human, X-ray, redundancy), not containing any of the previously selected ligands was used as negative data. All pockets were computed using fpocket (19) and all secondary structure information with STRIDE (41).

The ‘true’ pockets containing ligands were identified using euclidean distance. For each pocket in each structure, if any voronoi vertices from a pocket and any ligand atom distance is inferior or equal to 1 angstrom, then the pockets are labelled as ‘true’. All other pockets in the PDB file are by default categorized as ‘false’.

To train the predictive model PickPocket offers the possibility to use random forest, support vector machine algorithms or neural networks implemented in Python. The neural network is configured to test multiple architectures and automatically select the model with the best accuracy. The Python program included the following libraries: scikit-learn(41), numpy (42) and pandas (43).

Once the desired model it can be saved and used to make predictions on unseen data. A file containing all the Protein Data Bank pockets (all pockets from all structures) can be found in the data attached to the software.

